# Blockade of insulin-like growth factors increases efficacy of paclitaxel in metastatic breast cancer

**DOI:** 10.1101/165068

**Authors:** Lucy Ireland, Almudena Santos, Fiona Campbell, Carlos Figueiredo, Lesley Ellies, Ulrike Weyer-Czernilofsky, Thomas Bogenrieder, Michael Schmid, Ainhoa Mielgo

## Abstract

Breast cancer remains the leading cause of cancer death in women due to metastasis and the development of resistance to established therapies. Macrophages are the most abundant immune cells in the breast tumor microenvironment and can both inhibit and support cancer progression. Thus, gaining a better understanding of how macrophages support cancer could lead to the development of more effective therapies. In this study, we find that breast cancer associated macrophages express high levels of insulin-like growth factors 1 and 2 (IGFs) and are the main source of IGFs within both primary and metastatic tumours. 75% of breast cancer patients show activation of Insulin/IGF-1 receptor signaling and this correlates with increased macrophage infiltration and advanced tumor stage. In patients with invasive breast cancer, activation of Insulin/IGF-1 receptors increased to 87%. Blocking IGF in combination with paclitaxel, a chemotherapeutic agent commonly used to treat breast cancer, showed a significant reduction in tumor cell proliferation and lung metastasis in a pre-clinical breast cancer model compared to paclitaxel monotherapy. Our findings provide the rationale for further developing the combination of paclitaxel with IGF blockers for the treatment of invasive breast cancer, and Insulin/IGF1R activation and IGF+ stroma cells as potential biomarker candidates for further evaluation.

## INTRODUCTION

Breast cancer is the leading cause of cancer death in females worldwide, and is characterized by a high proliferation rate, an increased capacity to metastasize, and its ability to resist standard therapies (1). Triple negative breast cancer (TNBC) is a highly metastatic subtype of breast cancer that accounts for approximately 20% of all breast cancer cases and has limited efficacious treatment options (2). Current standard treatments for metastatic disease include radiotherapy and chemotherapy (3, 4). TNBC has a poorer survival rate, its biology is comparatively less well-understood and currently no effective specific targeted therapy is readily available (5). Breast cancer has a propensity to give rise to distant metastasis at sites such as the lungs, bone and brain, which can present up to ten years after treatment (6). Patients with metastatic breast cancer ultimately often become resistant to current chemotherapy treatments and as a result account for >90% of breast cancer deaths (7), highlighting the need for new therapeutic targets to treat metastatic burden more effectively.

Tumor progression and response to therapy is not only dependent on tumor intrinsic mechanisms but also involves modulation by surrounding non-malignant stromal cells in the tumor microenvironment (8, 9). Macrophages are the most abundant leukocytes in the breast tumor microenvironment (10) and an increase in tumor associated macrophages (TAMs) correlates with a poorer prognosis in patients (11–13). Macrophages can be polarized into M1-like anti-tumorigenic macrophages and M2-like pro-tumorigenic macrophages (14–16). M2-like macrophages can influence tumor initiation, progression, metastasis (17–19) and resistance to therapies (20–22). Cancer progression relies on the continued propagation of cancer cells which can be stimulated by external ligands activating signaling pathways of tumor cell survival and proliferation, even when challenged with chemotherapy (23–26). The insulin-like growth factor (IGF) signaling axis has been implicated in promoting cancer progression in several tumor types including breast cancer (27–29), and in breast cancer resistance to estrogen and HER2 receptor inhibition (27, 30–32).

Interestingly, Fagan et al., showed that tamoxifen resistant ER+ cells showed a reduction in the number of IGF-1 receptors while the number of insulin receptors and AKT phosphorylation levels remained unaltered when stimulated with Insulin and IGF-2, suggesting that both IGF-1 and IGF-2 signaling may support resistance of breast cancer cells to therapies (33). However, the role of IGF signaling in tumor progression and resistance to chemotherapy in breast cancer is not completely understood yet (32). We and others have recently shown that stroma-derived IGFs promote survival of cancer cells leading to therapy resistance in pancreatic and brain cancer models, respectively (22, 34). In the current studies, we aimed to investigate the role of stroma-derived IGF in breast cancer progression and metastasis, and to explore the therapeutic opportunity of blocking IGF signaling in combination with chemotherapy for the treatment of breast cancer.

## RESULTS

### Insulin and IGF-1 receptors are activated on tumor cells in biopsies from breast cancer patients, and this positively correlates with increased TAM infiltration and advanced tumor stage

Macrophages play an important role in breast cancer progression and metastasis (35, 36) and have been shown to express high levels of IGFs in other cancer types (22, 34), but the role of IGF-expressing macrophages in breast cancer has not yet been explored. To investigate whether IGF signaling pathways are activated in invasive breast cancer progression and whether their activation correlates with macrophage infiltration, we first evaluated the activation status of insulin and IGF-1 receptors in biopsies from breast cancer patients, and the levels of infiltrated tumor associated macrophage (TAMs). Immunohistochemical staining of serial sections of non-malignant breast tissue (Fig. 1A) and breast cancer patients’ tissues (Fig. 1B and C) revealed an increase in phospho-insulin/IGF-1 receptor levels in the breast cancer tissues along with increased infiltration of CD68+ (pan-myeloid/macrophage marker) and CD163+ (marker commonly used to identify pro-tumorigenic M2-like macrophages) macrophages (Fig. 1 A-D). Analysis of a tissue microarray (TMA) containing samples from 75 breast cancer patients, with different tumor stages but unspecified subtype, showed that Insulin/IGF-1R signaling was activated in 56 of 75 (~75%) patients (Fig. 1E and Supplementary Table. S1). Activation of insulin/IGF-1 receptors positively correlates with increased infiltration of CD163+ macrophages in the tumor (Fig. 1F) and with advanced tumor stage (Fig.1G).

**Figure 1.**
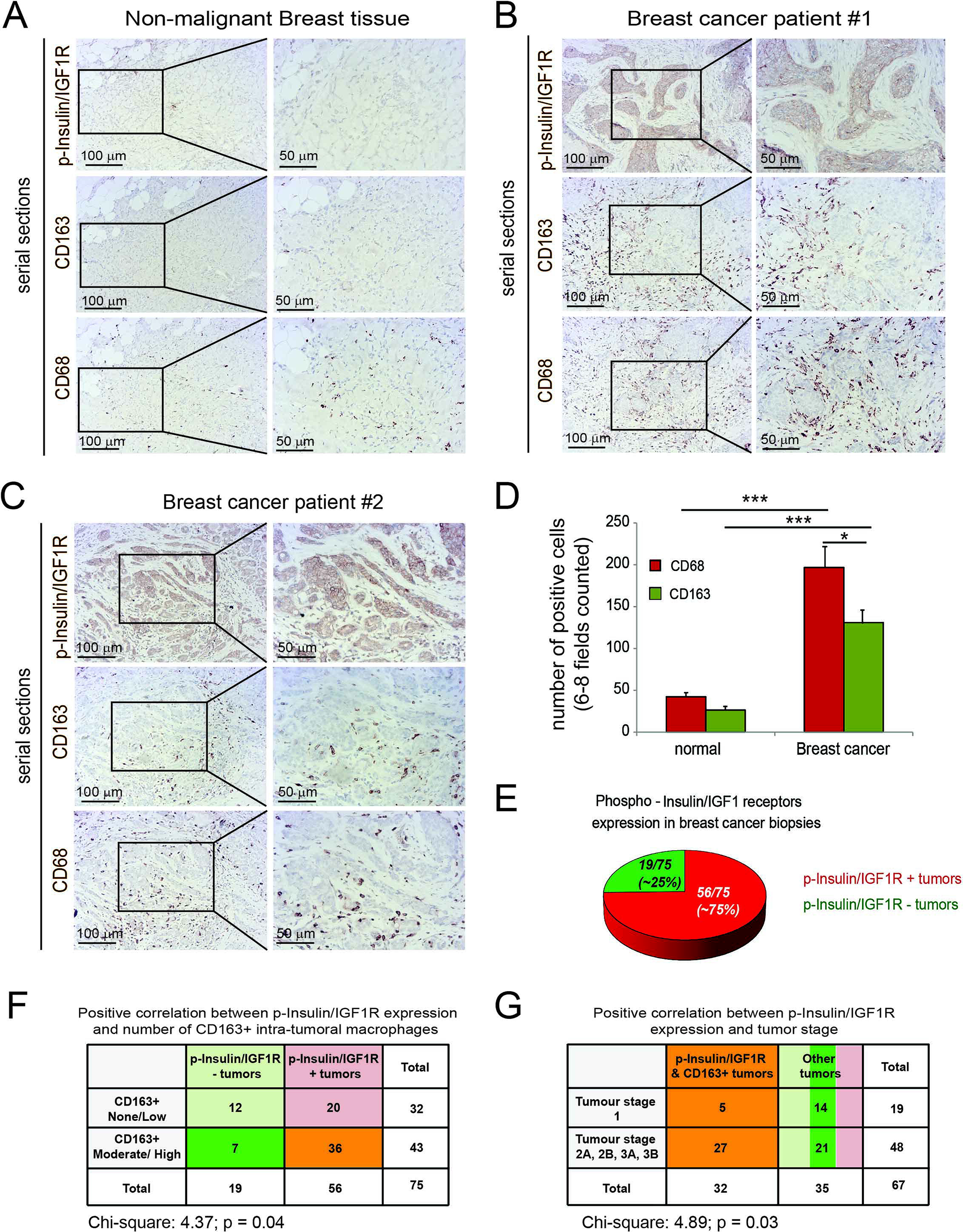
75% of breast cancer patients have activated Insulin/IGF1 receptors and Insulin/IGF-1 receptor activation positively correlates with macrophage infiltration and advanced tumor stage. **(A)** Serial sections of biopsies from non-malignant breast tissue immunohistochemically stained for phospho-Insulin/IGF1 receptor, CD163 and CD68. Scale bars, 100µm and 50µm. **(B) and (C)** Serial sections of biopsies from breast cancer patients immunohistochemically stained for phospho-Insulin/IGF1 receptor, CD163 and CD68. Scale bars, 100µm and 50µm. **(D)** Bar graph depicting the quantification of CD68 and CD163 positive macrophages in non-malignant breast tissue and breast cancer tissue samples. Error bars represent s.d. (n=3); * two tailed p value ≤ 0.05, *** two tailed p value ≤ 0.005 using a student's t-test. **(E)** Pie diagram representing the percentage of phospho-Insulin/IGF-1 receptor positive (red) and negative (green) tumors assessed in a tissue microarray containing biopsies from 75 breast cancer patients. **(F)** Contingency table and results from statistical analysis showing a positive correlation between phospho-Insulin/IGF-1 receptor expression in breast tumors and increased CD163+ macrophage infiltration. Chi-square = 4.37; p = 0.04. **(G)** Contingency table and results from statistical analysis showing a positive correlation between phospho-Insulin/IGF1 receptor and CD163+ macrophages co-expression and tumor stage. Chi-square = 4.89; p = 0.03.

### 87% of patients with invasive breast cancer have Insulin/IGF-1 receptors activated

Since Insulin/IGF-1 receptors activation positively correlates with advanced tumor stage, we further analysed biopsies from patients with invasive breast cancer. Immunohistochemical staining revealed that invasive breast cancer also shows increased phospho-insulin/IGF-1 receptor levels in tumor cells surrounded by CD163+ macrophages (Fig. 2A). Analysis of a TMA containing 90 samples from patients with invasive breast cancer showed that 78 of 90 (~87%) of these patients have Insulin/IGF1 receptors activated (Fig. 2B, upper pie diagram, and Supplementary Table S2). Among these 90 samples, 51 were TNBC of which 45 (~88.2%) showed activation of Insulin/IGF1 receptors (Fig. 2B, lower, left pie diagram), 13 were hormone receptor positive (HR+) of which 12 (~92%) showed activation of Insulin/IGF1 receptors (Fig. 2B, lower, middle pie diagram), and 19 were HER2 positive (HER2+) of which 16 (~84%) showed activation of Insulin/IGF1 receptors. Together these results suggest an important role for IGF signaling in invasive breast cancer of all subtypes, including TNBC.

**Figure 2.**
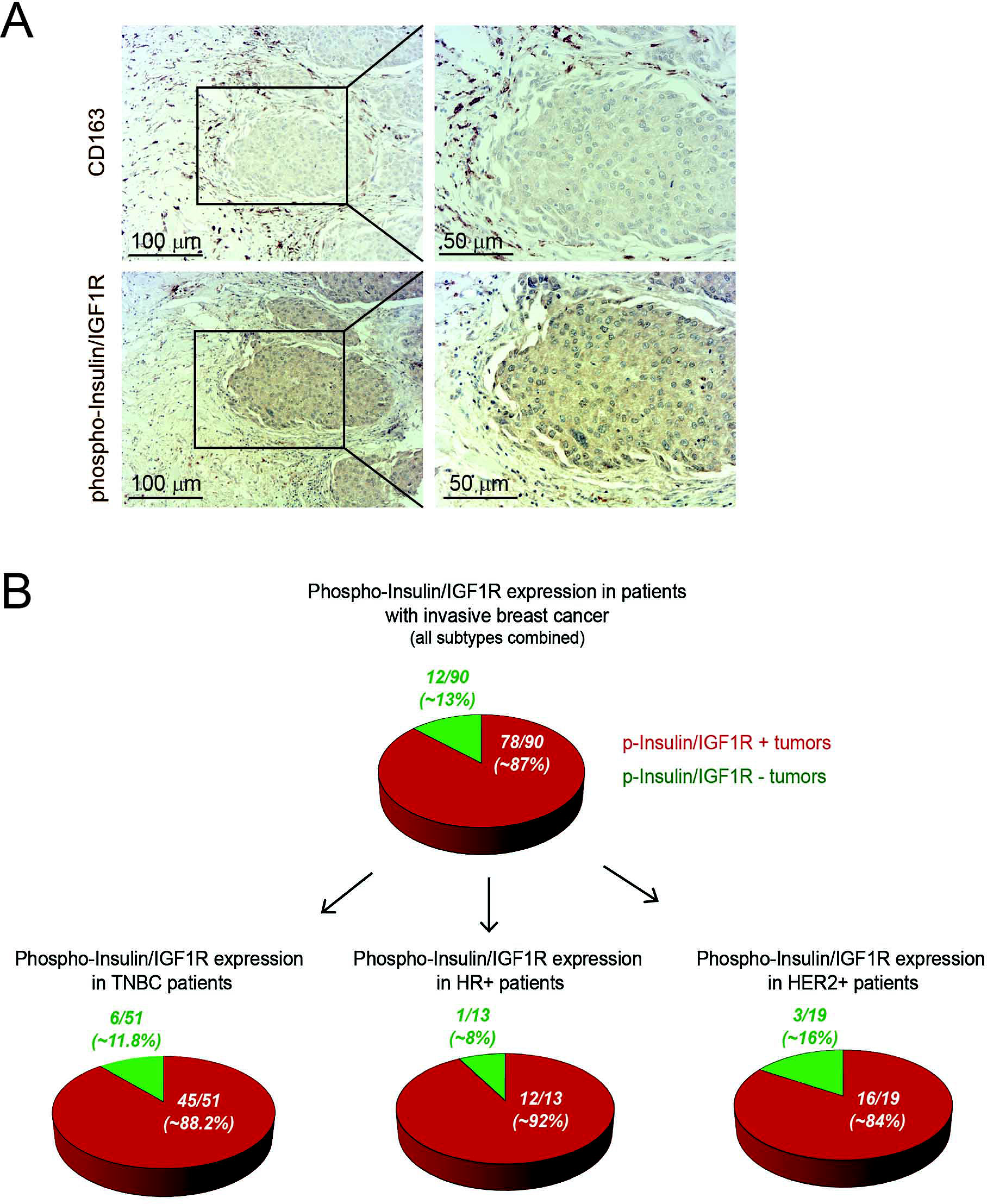
87% of patients with invasive breast cancer have activated Insulin/IGF1 receptors. **(A)** Immunohistochemical staining of invasive breast cancer tissue serial sections stained for phospho-Insulin/IGF1 receptor and CD163. Scale bars, 100 μm and 50 μm. **(B)** Upper diagram: Pie diagram representing the percentage of phospho-Insulin/IGF1 receptor positive (red) and negative (green) tumors assessed in tissue microarray (TMA) containing biopsies from 90 consented patients with invasive breast cancer. Lower diagrams: represent the percentage of phosphor-Insulin/IGF1 receptor positive (red) and negative (green) tumors of the molecular subsets, TNBC, HR+ and HER2+.

### Tumor associated macrophages and fibroblasts are the main sources of IGF-1 and IGF-2 in invasive breast cancer

To further understand the correlation between activation of Insulin/IGF1 receptors and increased TAMs infiltration, we used an orthotopic syngeneic TNBC pre-clinical model which has been shown to recapitulate the human disease progression (37). Briefly, we isolated murine breast cancer cells (Py230) from the genetically engineered spontaneous breast cancer model MMTV-PyMT and transduced isolated cells with a reporter lentivirus expressing zsGreen/luciferase allowing *in vivo* imaging of tumor burden. To induce breast tumor burden, Py230zsGreen/luciferase cells were orthotopically implanted into the mammary fat pad of isogenic immunocompetent recipient mice. 42 days after implantation, primary tumors were surgically removed and tumor tissue sections were analysed by immunofluorescent and immunohistochemical staining. We found that tumors were highly infiltrated by macrophages (F4/80+, CD68+, CD206+) that surround proliferating tumor cells (Ki67+) (Fig. 3A), which also express phosphorylated/activated insulin/IGF-1 receptors (Fig. 3B). Thus, similar to what we observed in human biopsies, murine breast cancer cells, which are spatially located in close proximity to M2-like TAMs, have activated Insulin/IGF-1 receptors.

**Figure 3.**
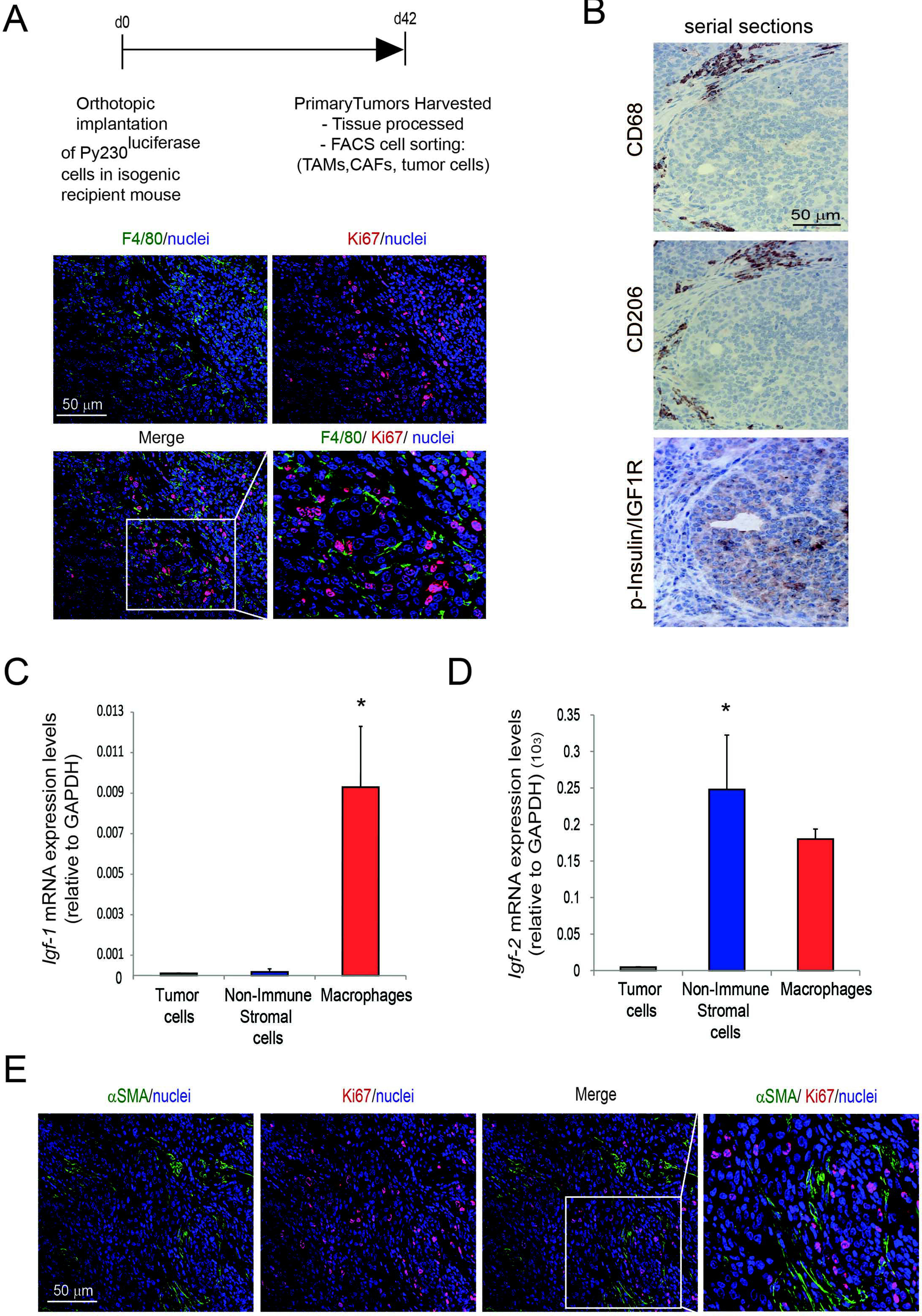
TAMs and CAFs are the main sources of IGF-1 and IGF-2 in invasive breast adenocarcinomas. **(A)** PY230 tumor cells were subcutaneously implanted into the third mammary gland of syngeneic recipient mice. Images show immunofluorescent staining for F4/80 (green), Ki67 (red) and nuclei (blue) in murine breast cancer tissue harvested at day 42 after tumor implantation. Scale bar 50 μm. **(B)** Serial sections of immunohistochemical staining for CD68, CD206 and phospho-Insulin/IGF1 receptor in murine breast tumors. Scale bar 50 μm. **(C)** *Igf-1* mRNA expression levels were quantified in tumor cells, non-immune stromal cells and tumor associated macrophages isolated from murine breast cancer tumors by flow cytometry. Error bars represent s.e. (n=3), * p value ≤ 0.05 using one-way ANOVA and Bonferroni *post-hoc* test. **(D)** *Igf-2* mRNA expression levels were quantified in tumor cells, non-immune stromal cells and tumor associated macrophages isolated from murine breast cancer tumors by flow cytometry. Error bars represent s.e. (n=3), * p value ≤ 0.05 using one-way ANOVA and Bonferroni *post-hoc* test. **(E)** Immunofluorescent images of αSMA (green), Ki67 (red) and nuclei (blue) in breast tumors. Scale bar 50 μm.

Next, we aimed to identify the source of insulin and IGF-1 receptors ligands, namely IGF-1 and IGF-2, in the tumor microenvironment. Therefore, we enzymatically digested primary tumors to prepare single cells suspensions and isolated tumor cells, non-immune stromal cells and macrophages, by flow cytometry cell sorting (Supplementary Fig.S1A,B). Gene expression analysis of isolated cell populations revealed that TAMs are the main source of *Igf-1* and that both TAMs and cancer associated fibroblasts (CAFs) are major sources of *Igf-*2 in the breast tumor microenvironment (Fig. 3C and D). Immunofluorescent staining of αSMA and Ki67 in the murine breast tumors showed that αSMA+ stromal cells, which appear to be a major source of IGF-2, also surround actively dividing tumor cells (Fig. 3E). IGF-1 and IGF-2 are also expressed in the stroma surrounding cancer cells in human TNBC samples (Supplementary Fig. S2A), and gene expression analysis of *Igf-1* and *Igf-2* in human MDA-MB-231 TNBC cells and primary human macrophages revealed that human macrophages express high levels of *Igf-1* and *Igf-2*, while there was very little expression of these ligands in the breast cancer cells (Supplementary Fig. S2B and C).

### Metastasis associated macrophages and fibroblasts remain the main sources of IGF-1 and IGF-2 in pulmonary metastatic lesions

Breast cancer is a highly invasive disease and often metastasizes to the lung. Orthotopically implanted Py230 breast cancer cells into syngeneic recipient mice effectively metastasized to the lung, where they formed metastatic tumors (Fig. 4A). We wondered whether IGF-1 and IGF-2 might also be expressed in the metastatic tumor microenvironment and thereby provide a survival/proliferative signal to disseminated cancer cells. To address this question, we analysed metastatic lung tumors from our animal model for the expression of *Igf-*1 and *Igf-*2 in the metastasis associated stromal cells.

**Figure 4.**
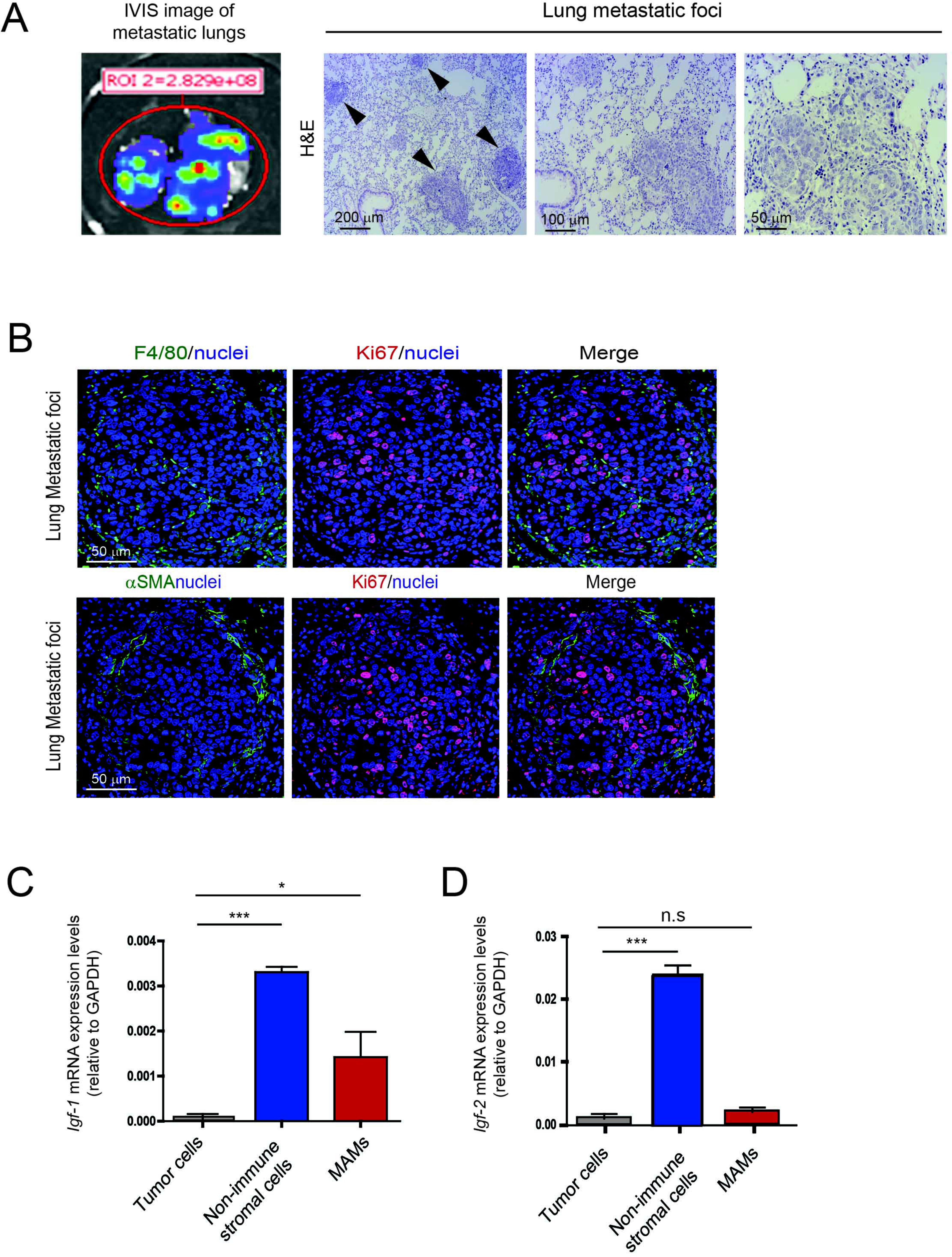
Metastasis associated macrophages and fibroblasts express IGF-1 and IGF-2 in metastatic lungs. **(A)** Left, identification of metastatic tumor lesions in the lung by bioluminescent imaging technique of orthotopically implanted PY230^luc^ breast cancer cells. Right, images show H&E staining of metastatic foci in the lungs. Arrows indicate metastatic foci, scale bars 200 μm, 100μm and 50 μm. **(B)** Immunofluorescent images of lung metastatic foci stained for F4/80 (green),αSMA (green), Ki67 (red) and nuclei (blue). Scale bar 50 μm. **(C)** Quantification of *Igf-1* mRNA expression levels in metastatic tumor cells, metastasis associated non-immune stromal cells and metastasis associated macrophages isolated from pulmonary metastasis. Error bars represent s.e. (n=3), * p value ≤ 0.05, *** p value ≤ 0.0001 using one-way ANOVA and Bonferroni *post-hoc* test. **(D)** Quantification of *Igf-2* mRNA expression levels in metastatic tumor cells, metastasis associated non-immune stromal cells and metastasis associated macrophages isolated from pulmonary metastasis. Error bars represent s.e. (n=3), ***p value ≤ 0.05 using one-way ANOVA and Bonferroni *post-hoc* test.

Metastatic tumors in lungs were first confirmed by bioluminescence *ex vivo* imaging and by hematoxylin and eosin (H&E) staining (Fig. 4A). immunofluorescent staining showed that pulmonary metastatic lesions are surrounded by macrophages (CD68+, F4/80+) and myofibroblasts (αSMA+) (Fig. 4B). Similar to what we observe at the primary site, metastasis associated macrophages (MAMs) and fibroblasts express high levels of *Igf-1* and *Igf-2*, while disseminated breast cancer cells do not express these ligands (Fig. 4C and D). Together, these findings provide evidence that macrophages and fibroblasts are the main sources of IGF-1 and IGF-2 both at the primary and the metastatic site in invasive breast cancer.

### Combination treatment of invasive breast cancer with paclitaxel and IGF blocking antibody reduces tumor cell proliferation and metastasis

To determine whether IGF signaling affects breast cancer progression, metastasis and response to paclitaxel, a standard chemotherapeutic agent used to treat breast cancer, we treated mice orthotopically implanted with Py230 TNBC cells, with control IgG antibody, IGF-1/2 blocking antibody xentuzumab, paclitaxel or xentuzumab with paclitaxel (Fig. 5A). As expected, control and paclitaxel treated mice showed high levels of insulin and IGF-1 receptor activation in the primary breast cancer tumors, whereas the xentuzumab and xentuzumab with paclitaxel treated groups showed markedly reduced levels of insulin and IGF-1 receptor activation, confirming that xentuzumab has reached the tumor and has blocked IGF signaling (Fig. 5B) (38). No differences were seen in primary tumor growth (Supplementary Fig. S3A), in tumor cell death (Supplementary Fig. S3B) or in TAM infiltration of primary tumors (Supplementary Fig. S3C) between the different treatment groups. However, control IgG treated mice showed higher levels of Ki67+ proliferating tumor cells which were modestly reduced by both paclitaxel and xentuzumab single treatments and significantly reduced by the combination treatment of xentuzumab with paclitaxel (Fig. 5C and D). In addition, mice treated with the combination of xentuzumab with paclitaxel treatment showed a reduction in lung metastasis incidence (Fig. 5E). Although there were no significant differences in the number of metastatic foci in the different treatment groups (Fig. 5F), we found a significant reduction in the size of metastatic lesions in the group treated with both paclitaxel and xentuzumab (Fig. 5G and H). These data suggest that initial metastatic seeding is not affected by the combination treatment, but that paclitaxel/xentuzumab treatment impairs metastatic outgrowth of disseminated breast cancer cells.

**Figure 5.**
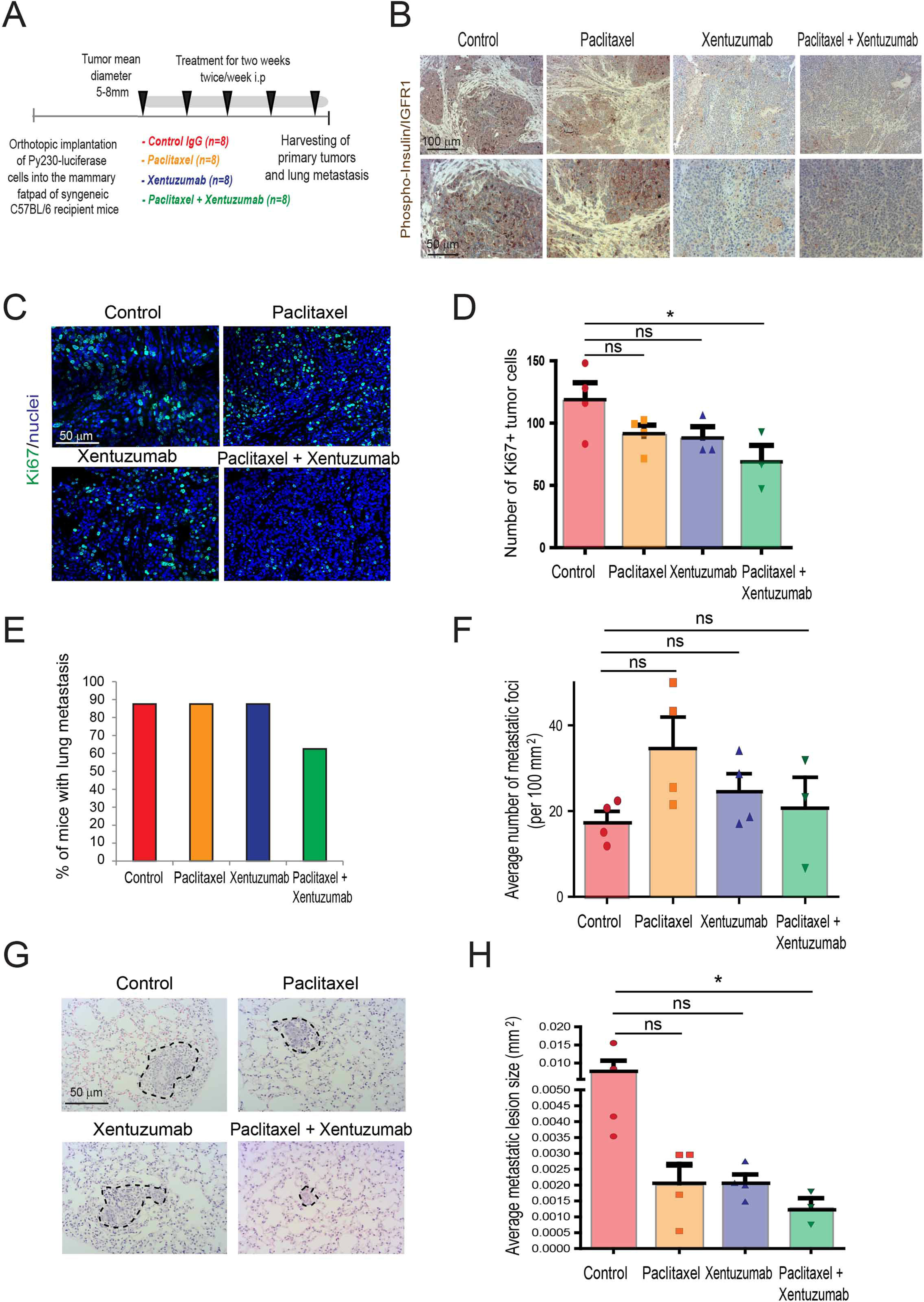
Combined treatment of IGF blocking antibody with paclitaxel decreases breast cancer proliferation and metastasis. **(A)** Py230-luciferase cells were orthotopically implanted into the third mammary fatpad of syngeneic C57BL/6 recipient mice and mice were treated, starting when tumors reached between 5-8mm^2^, twice a week i.p., with control IgG antibody, IGF blocking antibody xentuzumab (100mg/kg), paclitaxel (100mg/kg) or a combination of xentuzumab with paclitaxel (n = 8 mice/group). **(B)** Immunohistochemical staining of phospho-insulin/IGF-1R in breast tumors treated with IgG (control), paclitaxel, xentuzumab or paclitaxel + xentuzumab. Scale bars 100 μm and 50 μm. **(C)** Immunofluorescent staining of Ki67 in primary tumors treated with IgG (control), paclitaxel, xentuzumab or paclitaxel + xentuzumab. Scale bar 50 μm. **(D)** Quantification of Ki67 positive tumor cells in tumors treated with IgG (control), paclitaxel, xentuzumab or paclitaxel + xentuzumab. 3-5 fields counted/ mouse tumor, n=3-4 mice per treatment group, * p ≤ 0.05 using one-way ANOVA and Bonferroni *post-hoc* test. **(E)** Percentage of mice presenting with lung metastasis per treatment group. (n=8 mice/group). **(F)** Quantification of number of lung metastatic foci per 100mm2 in mice treated with control IgG, paclitaxel, xentuzumab or paclitaxel + xentuzumab. ns, non-significant differences using one way ANOVA and Bonferroni *post-hoc* test. **(G)** H&E staining of lung metastatic foci in mice treated with control IgG, paclitaxel, xentuzumab or paclitaxel + xentuzumab. Scale bar 50 μm. **(H)** Average size of pulmonary metastatic lesions in mice treated with control IgG, paclitaxel, xentuzumab or paclitaxel + xentuzumab, * p ≤ 0.05, using one-way ANOVA and Bonferroni *post-hoc* test.

Taken together, our findings indicate that IGF-1 and 2 are highly expressed by both macrophages and fibroblasts in invasive breast cancer, and that blockade of IGF potentiates the efficacy of paclitaxel.

## DISCUSSION

Here we show that IGF-1 and 2 are secreted by macrophages and fibroblasts, both at the primary site and metastatic site in invasive breast cancer, and that blocking IGF increases the efficacy of paclitaxel, a chemotherapeutic agent commonly used for the treatment of invasive breast cancer (Fig. 6). Breast cancer, and in particular TNBC, remains a highly metastatic and potentially lethal disease with a need to identify additional specific molecular targets and to develop more effective therapies (4, 5). While IGF signaling has been shown to support progression of HR+ and HER2+ breast cancer and the development of resistance to established therapies, its precise role in TNBC remains elusive (32). In our TNBC model, we found that TAMs and CAFs secrete IGF-1 and 2 at both the primary site and the pulmonary metastatic site. Zhang et al., previously reported that CAF-derived IGF-1 primes breast cancer cells for bone metastasis (39). These studies both suggest that stromal-derived IGF plays an important role in the metastatic process of breast cancer.

**Figure 6.**
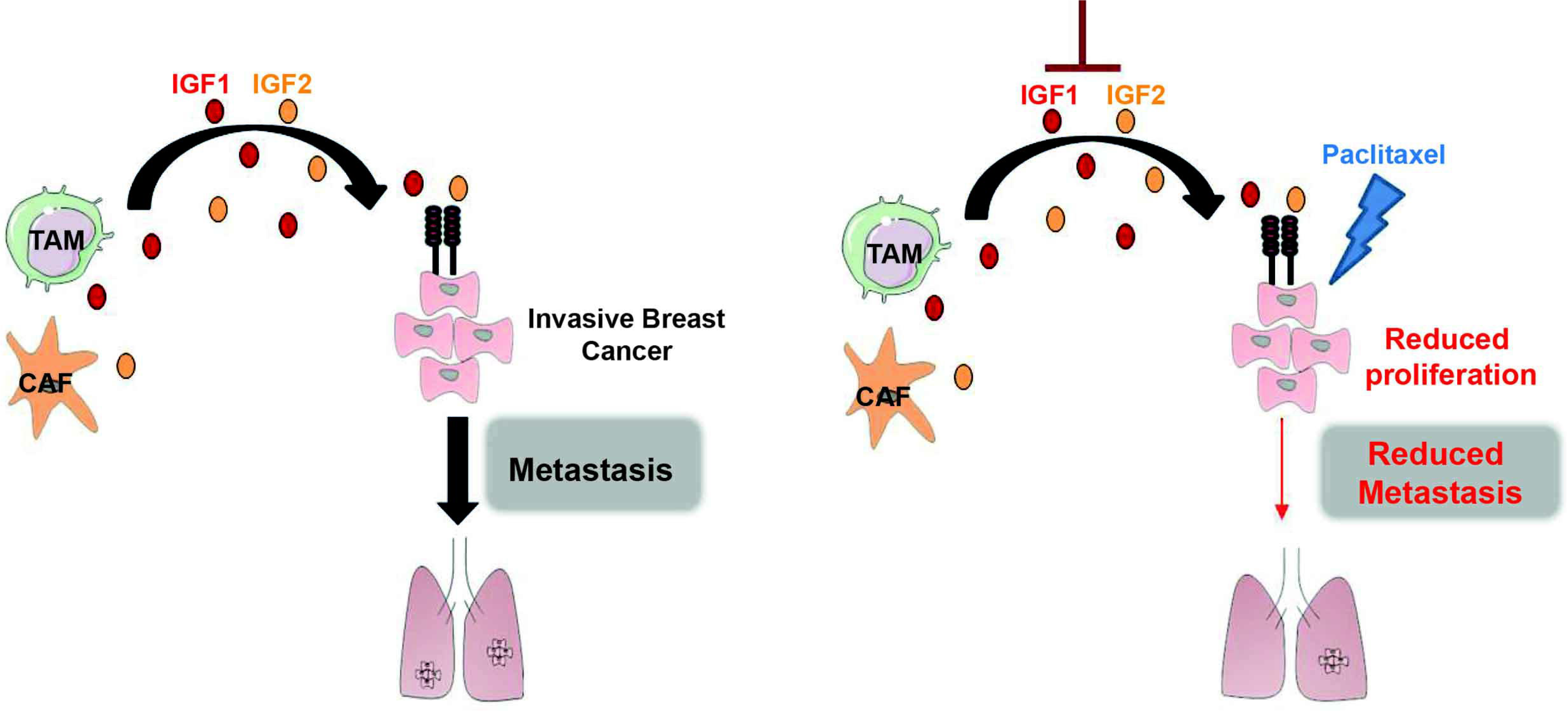
Combined treatment of IGF blocking antibody xentuzumab with paclitaxel decreases breast cancer metastasis. Schematics describing the role of stroma-derived IGF-1 and 2 in regulating the response of metastatic breast cancer to paclitaxel.

In this study, we find that the Insulin/IGF1R signaling pathway is activated in 78 of 90 (~87%) of patients with invasive breast cancer and 45 of 51 (~88.2%) of TNBC patients, suggesting IGF may be a promising therapeutic target for this highly aggressive breast cancer subtype. We also observed that activation of Insulin/IGF1 receptor signaling positively correlates with increased levels of macrophage infiltration and advanced tumor stage in patients, suggesting that Insulin/IGF1 receptor activation and/or stroma expression of IGF could be predictive biomarker candidates for further evaluation.

To investigate the therapeutic potential of blocking IGF signaling in invasive breast cancer, we tested the IGF-1/2 blocking antibody xentuzumab (Boehringer Ingelheim) in a pre-clinical mouse model of invasive TNBC breast cancer which metastasizes to the lungs. We found that the combination of paclitaxel with xentuzumab significantly reduced tumor cell proliferation, frequency of lung metastasis and size of metastatic lesions. In agreement with our findings, Gooch et al., previously showed that IGF-1 promotes proliferation of paclitaxel treated cells *in vitro* (40).

IGF1R inhibitors have been assessed in clinical trials for metastatic hormone receptor positive (HR+) as well as triple negative breast cancer but have shown limited success (41–46). Two promising IGF ligand blocking antibodies, xentuzumab and MEDI-573, are currently being evaluated in clinical trials in HR+ metastatic breast cancer patients in combination with everolimus and exemestane (NCT02123823) and in hormone sensitive metastatic breast cancer in combination with letrozole (NCT01446159) respectively. In contrast to IGF1R antibodies, IGF blocking antibodies neutralize both ligands IGF-1 and IGF-2 and thereby inhibit proliferative signaling through both Insulin and IGF1 receptors without affecting insulin metabolic signaling (38, 47).

Breast cancer cells survive poorly in isolation and participate in a complex relationship with surrounding stromal and immune cells in the tumor microenvironment which can support tumor cell survival, proliferation and spreading to other organs (17, 20, 21, 36, 39). CAFs and TAMs are the most abundant stromal cells in solid cancers, including breast cancer. However, different populations of CAFs and TAMs with both pro- and anti-tumorigenic functions co-exist in tumors (48–50). Therefore, therapies aiming to specifically inhibit the tumor supporting functions of stromal cells, without affecting their anti-tumorigenic functions, may be more effective than ablation therapies in restraining tumor progression (26, 51). Our findings indicate that blocking IGFs in combination with paclitaxel, decreases tumor cell proliferation and breast cancer pulmonary metastasis without affecting macrophage infiltration. In conclusion, this study suggests that stroma-derived IGFs support breast cancer metastasis and modulate its response to paclitaxel, providing the rationale for further evaluation of IGF blocking antibodies in combination with paclitaxel in the treatment of invasive breast cancer.

## Materials and Methods

### Generation of primary PyMT-derived breast cancer cells

Py230 cells (hormone receptor negative and HER2 low) were generated in Ellies lab (University of California San Diego, U.S.A) and obtained from spontaneously arising tumors in MMTV-PyMT C57Bl/6 female mice by serial trypsinization and limiting dilution (52). The mouse model used for obtaining these tumors has been described in detail previously (53, 54).

### Cell lines and culture conditions

Murine Py230 cells were cultured in DMEM/F-12 culture media supplemented with 10% FBS and supplemented with MITO serum extender (Corning #355006), 1% penicillin/streptomycin at 37°C, in a 5% CO_2_ incubator. Human MDA-MB231 breast cancer cells were cultured in DMEM supplemented with 10% FBS, 1% penicillin/streptomycin, at 37°C, 5% CO_2_ incubator. Cells were authenticated, and periodically tested for mycoplasma contamination.

### Generation of primary macrophages

Primary human macrophages were generated by purifying CD14+ monocytes from blood samples obtained from healthy subjects using magnetic bead affinity chromatography according to manufacturer's directions (Miltenyi Biotec) followed by incubation for 5 days in RPMI containing 10% FBS and 50 ng/mL recombinant human M-CSF (Peprotech).

### Syngeneic orthotopic breast cancer model

2 × 10^6^ Py230 luciferase/zsGreen labelled cells were injected into the fat pad of the third mammary gland of C57BL/6 6-8 week-old female mice. Tumors were measured with calipers twice a week and treatment was started when tumors started to grow and measured between 5-8mm mean diameter. Mice were administered i.p with IgG isotype control antibody, paclitaxel (100 mg/kg), IGF-1/2 blocking antibody xentuzumab (100 mg/Kg) (38) kindly provided by Boehringer Ingelheim, or Paclitaxel with xentuzumab, twice a week for 15 days. At humane endpoint, primary tumors and lungs were harvested, imaged using IVIS technology and tissues were either digested for FACS sorting and analysis (see details below) or formalin fixed and paraffin embedded. Paraffin embedded lungs were serially sectioned through the entire lung. Sections were stained with hematoxylin and eosin (H&E), and images were taken using a Zeiss Observer Z1 Microscope. Number of foci and total area of metastatic foci were calculated to estimate seeding and metastasis burden using ZEN imaging software.

### FACS sorting and analysis of tumors

Single cell suspensions from murine primary breast tumors and pulmonary metastasis were prepared by mechanical and enzymatic disruption in Hanks Balanced Salt Solution (HBSS) with 1 mg/mL Collagenase P (Roche). Tumor cells, tumor associated macrophages and stromal cells were analysed and sorted using flow cytometry (FACS ARIA II, BD Bioscience). Cell suspensions were centrifuged for 5 min at 1500 rpm, resuspended in HBSS and filtered through a 500 μm polypropylene mesh (Spectrum Laboratories). Cells were resuspended in 1 mL 0.05%Trypsin and incubated at 37°C for 5 minutes. Cells were filtered through a 70 μm cell strainer and resuspended in PBS + 1% BSA, blocked for 10 minutes on ice with FC Block (BD Pharmingen, Clone 2.4G2) and stained with Sytox® blue viability marker (Life Technologies) and conjugated antibodies against anti-CD45-PE/Cy7 (Biolegend, clone 30-F11) and anti-F4/80-APC (Biolegend, clone BM8). Cell analysis was performed using FACS Canto II.

### Gene expression

Total RNA was isolated from FACS sorted tumor cells, tumor associated macrophages and stromal cells from primary breast tumors and lung metastasis as described in Qiagen Rneasy protocol. Total RNA from the different cell populations was extracted using a high salt lysis buffer (Guanidine thiocynate 5 M, sodium citrate uM, lauryl sarcosine 0.5% in H2O) to improve RNA quality followed by purification using Qiagen Rneasy protocol. cDNA was prepared from 1μg RNA/sample, and qPCR was performed using gene specific QuantiTect Primer Assay primers from Qiagen. Relative expression levels were normalized to *gapdh* expression according to the formula <2^− (Ct *gene of interest* – Ct *gapdh*) (55).

### Tissue Microarrays

A tissue microarray (TMA) containing 75 breast cancer samples from consented patients was purchased from the Liverpool Tissue Bank. This TMA did not provide information of cancer subtypes. A second tissue microarray BR10011 containing 90 invasive breast cancer samples was purchased from U.S Biomax. Among these 90 invasive breast cancer samples 51 were TNBC, 13 were hormone receptor positive (HR+) and 19 were HER2+. Both TMAs were subjected to immunohistochemical staining and scoring by a pathologist. Detailed information of both TMAs is provided in Supplementary Tables S1 and S2.

### Immunohistochemistry and Immunofluorescence

Deparaffinization and antigen retrieval was performed using an automated DAKO PT-link. Paraffin-embedded human and mouse breast tumors and lung metastasis were immuno-stained using the DAKO envision+ system-HRP.

#### Antibodies and procedure used for Immunohistochemistry

All primary antibodies were incubated for 2 hours at room temperature: CD68 (Human: DAKO, clone KP1, M081401-2 used 1:2000 after high pH antigen retrieval. Mouse: Abcam, ab31630 used at 1:100 after low pH retrieval), CD163 (Abcam, ab74604 pre-diluted after low pH antigen retrieval), Phospho-Insulin/IGF-1R (R&D, AF2507, used 1:50 after high pH antigen retrieval), phospho-insulin receptor (Lifespan Biosciences, LS-C177981 used 1:100 after low pH antigen retrieval), phospho-IGF-1R (Biorbyt, orb97626 used at 1:200 after high pH retrieval), IGF-1 (Abcam, ab9572 used at 1: 200 after high pH antigen retrieval), IGF-2 (Abcam, ab9574 used at 1: 200 after low pH antigen retrieval), CD206 (Abcam, ab8918 used 1:200 after low pH antigen retrieval). Subsequently, samples were incubated with secondary HRP-conjugated antibody (from DAKO envision kit) for 1 hour at room temperature. All antibodies were prepared in antibody diluent from Dako envision kit. Staining was developed using diamino-benzidine and counterstained with hematoxylin.

#### Antibodies and procedure used for Immunofluorescence on paraffin embedded tissues

Tissue sections were incubated overnight at RT with the following primary antibodies F4/80 (Biolegend, #123102 used at 1:200 after low pH antigen retrieval), Ki67 (Abcam ab15580 used at 1:1000 after low pH antigen retrieval) αSMA (Abcam ab7817 used at 1:100, after low pH antigen retrieval). Samples were washed with PBS and incubated with goat anti-rat 488 (Abcam ab96887), goat anti-rabbit 594 (Abcam ab98473) and goat anti mouse 488 (Abcam ab98637) secondary antibodies respectively all used at 1:300 and DAPI at 1:600 for 2 hours at RT. Slides were washed with PBS, final quick wash with distilled water and mounted using DAKO fluorescent mounting media.

### Statistical Methods

Statistical significance for *in vitro* assays and animal studies was assessed using unpaired two-tailed Student *t* test or one-way ANOVA coupled with Bonferroni's *post hoc* tests, and the GraphPad Prism 5 program. All error bars indicate SD of n=3 (*in vitro* studies) or SEM n=3-8 (animal studies). Human samples were analyzed using Fisher's exact test and the Matlab version 2006b program.

### Institutional approvals

All studies involving human tissues were approved by the University of Liverpool and were considered exempt according to national guidelines. Human breast cancer samples were obtained from the Liverpool Tissue Bank and patients consented to use the surplus material for research purposes or purchased from U.S Biomax. All animal experiments were performed in accordance with current UK legislation under an approved project licence (reference number: 403725). Mice were housed under specific pathogen-free conditions at the Biomedical Science Unit at the University of Liverpool.

All studies involving blood collection were approved by the National Research Ethics (Research Integrity and Governance Ethics committee-Reference: RETH000807). All individuals provided informed consent for blood donation on approved institutional protocols.

## AUTHOR CONTRIBUTIONS

L.I and A.S performed most of the experiments including immunohistochemical stainings, *in vivo* experiments, flow cytometry, cell isolations and qPCR. F.C. helped with the analysis and interpretation of tumor biopsies and tissue microarrays. C. F helped with FACS analysis. U.W and T.B provided with xentuzumab and technical advice for *in vivo* use of xentuzumab. L.E generated MMTV-PYMT-derived (Py230) primary breast cancer cells. M.C.S. provided conceptual advice and helped with *in vivo* experiments. A.M and L.I designed experiments and wrote the manuscript. A.M. conceived and supervised the project. All authors helped with the analysis and interpretation of the data, the preparation of the manuscript, and approved the manuscript.

## ACKNOWLEDGMENTS

These studies were supported by a Sir Henry Dale research fellowship to Dr Ainhoa Mielgo, jointly funded by the Wellcome Trust and the Royal Society (grant number 102521/Z/13/Z), a New Investigator research grant from the Medical Research Council to Dr Michael Schmid (grant number MR/L000512/1) and North West Cancer Research funding. Generation of primary breast cancer cells was supported by funding provided to Dr Lesley Ellies (K22CA118182).

We thank Dr. Ilaria Malanchi for reading the manuscript and providing critical feedback. We also thank Professor Azzam Taktak for assistance with biostatistical analysis. We thank Dr Arthur Taylor and Professor Patricia Murray for transducing the Py230 cells with zsGreen/luciferase lentivirus and Valeria Quaranta for assisting with FACS sorting. We acknowledge the Liverpool Tissue Bank for providing tissue samples, the flow cytometry/cell sorting facility, the biomedical science unit and the pre-clinical *in vivo* imaging facility for provision of equipment and technical assistance. We thank the patients and their families, as well as the healthy blood donors who contributed with tissue samples and blood donations to these studies.

